# Inhibition of Microbial Methane Oxidation by 2-Chloro-6-Methylpyridine

**DOI:** 10.1101/2022.10.13.512149

**Authors:** Edward J. O’Loughlin, Dionysios A. Antonopoulos, Kristin K. Arend, Theodore M. Flynn, Jason C. Koval, Sarah M. Owens

**Affiliations:** Biosciences Division, Argonne National Laboratory, Argonne, IL 60439-4843, USA; Old Woman Creek National Estuarine Research Reserve, Huron, OH 44839, USA

**Keywords:** methane oxidation, 2-chloro-6-methylpyridine, methanotrophs, nitrapyrin

## Abstract

Several pyridine derivatives including the pesticide nitrapyrin [2-chloro-6-(trichloromethyl) pyridine] are strong inhibitors of methane monooxygenase, a key enzyme of aerobic methane (CH_4_) oxidation. In this study we examined the effects of 2-chloro-6-methylpyridine (2C6MP) concentration on aerobic CH_4_ oxidation and the development of populations of putative methanotrophs in sediment from Old Woman Creek, a freshwater estuary in Huron Co., Ohio. Experimental systems were prepared in serum bottles containing minimal medium with a headspace containing 20% O_2_ and 10% CH_4_. The microcosms were spiked with 2C6MP to achieve concentrations of 0, 0.1, 1, or 10 mM and inoculated with sediment. When headspace CH_4_ concentrations decreased from 10% to < 2%, subsamples were taken for DNA extraction and sequencing of 16S rRNA gene amplicons. There was minimal effect of 2C6MP on CH_4_ oxidation at concentrations of 0.1, and 1 mM, but complete inhibition for > 20 months was observed at 10 mM. ANOSIM of weighted UniFrac distances between groups of triplicate samples supported a primary distinction of the inoculum relative to the enrichments (R=0.999) and a secondary distinction between bottles containing 2C6MP versus those without (R=0.464 [0.1 mM]; R=0.894 [1 mM]). The inoculum was dominated by members of the *Proteobacteria* (49.9±1.5%), and to a lesser extent by *Bacteroidetes* (8.8±0.2%), *Acidobacteria* (8.9±0.4%), and *Verrucomicrobia* (4.4±0.3%). In enrichments with or without 2C6MP, *Proteobacteria* expanded to comprise 65–70% of the total. In the absence of inhibitor, members of the *Methylococcaceae* and *Methylophilaceae* increased in relative abundance from < 0.1% of the inoculum to 8.5±1.0% and 13.4±2.3%, of the total community respectively. At both 0.1 and 1 mM concentrations of the inhibitor, the *Methylococcaceae* were much less abundant, representing 3.3±0.5% and 2.8±3.3% respectively. No inhibition of the *Methylophilaceae* was seen at the lower concentration of 2C6MP, but at the higher concentration this taxon was only 7.8±1.1% of the total. In contrast, members of the *Crenotrichaceae*, another group of methane oxidizers, increased in relative abundance with greater amounts of inhibitor, representing 8.6±3.6% of the total at 0.1 mM and 12.0±4.5% at 1 mM, compared to only 4.1±0.4% when no inhibitor was present. These results clearly show changes in the populations of putative aerobic methanotrophs relative to the amount of 2C6MP present.

## 1. Introduction

An improved understanding of the global terrestrial C cycle has become a policy imperative, both domestically and internationally, and is crucial in efforts to model, predict, and potentially mitigate the effects of increasing concentrations of greenhouse gasses on global climate. Although not as prevalent in the atmosphere as carbon dioxide (CO_2_), methane (CH_4_) is an important greenhouse gas that accounts for ∼20% of human-induced radiative forcing. Atmospheric CH_4_ concentrations have increased by nearly 160% since 1850, largely due to human activities relating to large-scale land management and agricultural practices (e.g., wetland rice production, raising of ruminant livestock, and mining operations). Moreover, increased CH_4_ emissions due to the warming of Arctic permafrost have been identified as a potentially significant factor resulting from (and contributing to) global climate change.

The formation (methanogenesis) and consumption of CH_4_ (methane oxidation) in soils and sediments are the result of highly specialized microorganisms [1]. Methanogenesis is carried out by a group of archaea called methanogens, which are obligate anaerobes in the domain *Archaea*. In most anoxic environments (e.g., saturated soils, wetlands, lacustrine and marine sediments) methanogenesis is the final process in the anaerobic degradation of organic carbon. Most methanogens use CO_2_ as the terminal electron acceptor in anaerobic respiration, reducing it to CH_4_ (hydrogenotrophic methanogenesis) [2]. However, a limited number of methanogens can generate CH_4_ from acetate fermentation (acetoclastic methanogenesis) [3]. Methane formed during methanogenesis can be oxidized to carbon dioxide via aerobic CH_4_ oxidation by methanotrophic bacteria—obligate aerobes that use CH_4_ as a sole C and energy source. Aerobic methanotrophs are found in a diverse range of aquatic and terrestrial environments including lacustrine, estuarine, and marine waters/sediments and soils ranging from arctic to tropical regions, and many are adapted to environments with extremes in pH (1–11), temperature (0–72 °C), and salinity (up to 30%) [4]. Despite the wide range in habitats, aerobic methanotrophic bacteria are not broadly distributed taxonomically, clustering primarily into two groups that differ with respect to phylogeny, ultrastructure, lipid composition, biochemistry, and physiology; type I methanotrophs are found within the *Gammaproteobacteria* in the family *Methylococcaceae* and type II methanotrophs are found within the *Alphaproteobacteria* in the family *Methylocystaceae*. Recently, a third group of aerobic methanotrophs have been identified comprising a single phylogenetic subcluster within the phylum *Verrucomicrobia* [5]. In addition to their role in C cycling via CH_4_ oxidation, aerobic methanotrophs also play a role in N cycling via ammonium oxidation and N_2_ fixation.

The pesticide nitrapyrin [2-chloro-6-(trichloromethyl) pyridine] is used to minimize the loss of nitrogen fertilizer (as ammonium) from soil by inhibiting ammonia monooxygenase (AMO), a key enzyme in the oxidation of ammonia to nitrate by nitrifying bacteria. However, AMO shares many similarities with methane monooxygenase (pMMO), a key enzyme of aerobic CH_4_ oxidation, and nitrapyrin has been shown to directly inhibit aerobic CH_4_ oxidation [6-8]. In this study we examined the effects of the nitrapyrin analog 2-chloro-6-methylpyridine (2C6MP) (Figure 1) on aerobic CH_4_ oxidation and microbial community development in freshwater wetland sediment.

**Figure 1.**
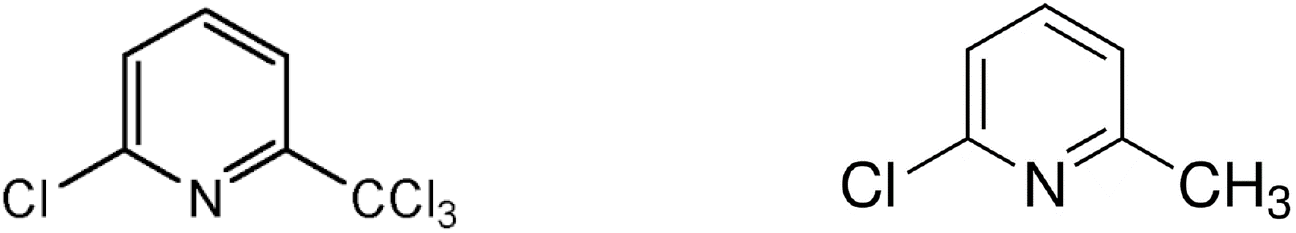
Molecular structure of nitrapyrin (left) and 2-chloro-6-methylpyridine (2C6MP) (right).

## 2. Materials and Methods

Overlying water and sediment (0-5 cm depth) were collected from Old Woman Creek, a freshwater estuary on the southern shore of Lake Erie in Huron Co., Ohio (41°22’35.472”N 82°30’29.999”W). The Old Woman Creek watershed encompasses 7,000 hectares, of which over 66% is agricultural land. The sediment and water were combined to create a slurry, that was used to inoculate the experimental systems described below.

A defined minimal medium (DMM) consisting of 20 mM PIPES buffer, 20 mM HEPES buffer, 1 mM NH_4_Cl, 1 mM KCl, 1 mM CaCl_2_, 1 mM MgCl_2_, 10 µM phosphate, 10 mL of trace metal solution [9], adjusted to pH 7.5, was sterilized by filtration through a 0.22 µm filter. The experimental systems were prepared in sterile 160-mL serum bottles containing 50 mL of DMM with a headspace consisting of 70% argon (Ar), 20% oxygen (O_2_), and 10% CH_4_, with 40 µmol of xenon (Xe) (as an internal standard), and sealed with butyl rubber plug stoppers and aluminum crimp caps. The microcosms were spiked with neat 2C6MP (99%, Sigma-Aldrich) to achieve concentrations of 0, 0.1, 1, or 10 mM (4 replicate microcosms at each concentration) and inoculated with 5 mL of sediment slurry. The microcosms were incubated at 30 ºC in the dark and monitored for evidence of CH_4_ oxidation using gas chromatography to measure changes in CH_4_ and O_2_ concentrations in the headspace over time. The concentrations of CH_4_, and O_2_ were measured by analyzing a 200 µL sample of headspace with an Agilent 7890A gas chromatograph equipped with a Supelco Carboxen 1010 Plot column (30 m × 0.35 mm) and a thermal conductivity detector, using Ar as the carrier gas (2.68 mL min^-1^) and a temperature program of 60–100 °C at 5 °C min^-1^, 100–230 °C at 40 °C min^-1^, and held at 230 °C for 3.4 min. The system was calibrated by equilibrating known masses of analyte (CH_4_ and O_2_) and internal standard (Xe) in serum bottles having the same ratio of aqueous phase to vapor phase as the experimental systems, thereby accounting for water-vapor partitioning. When headspace CH_4_ concentrations decreased from 10% to < 2%, a 10 mL sample of suspension was collected from each experimental system and frozen at −80 °C for subsequent DNA extraction and sequencing; samples for DNA extraction were not collected from the 10 mM 2C6MP treatments due to complete inhibition of CH_4_ oxidation.

Samples were mixed prior to removing 0.5 mL of suspension for transfer to extraction tubes. DNA extraction was then performed using the MOBIO PowerSoil DNA Isolation kit according to the manufacturer’s protocol. All DNA quantitation was performed using the Qubit assay (Invitrogen). The V4 region of the 16S rRNA gene (515F-806R) was PCR amplified as detailed in Caporaso et al. [10] to survey the total bacterial community in the extracted samples using the Earth Microbiome Project barcoded primer set adapted for the Illumina MiSeq https://earthmicrobiome.org/protocols-and-standards/16s/. Amplicons were then sequenced on a 151bp x 12bp x 151bp MiSeq run using customized sequencing primers and procedures as described by Caporaso et al. [11]. Triplicate libraries were prepared and sequenced for each sample as technical replicates. A total of 1.5 million paired sequences were generated (37,536 + 16,323 per amplicon library) and then merged followed by downstream processing using QIIME [12]. Briefly, singletons were removed, de novo OTU picking with uclust was used to cluster sequences at 97% similarity, and Greengenes (4feb2011) was used to assign taxonomies. For analyses of beta diversity, the total number of sequences in each library was normalized to the amount in the library containing the least number of sequences (7,168).

## 3. Results and Discussion

### 3.1. Inhibition of Methane Oxidation by 2C6MP

In the absence of 2C6MP, >80% of the CH_4_ was oxidized within 11 days (Figure 2). The presence of 0.1 or 1 mM 2C6MP resulted in a limited inhibition of CH_4_ consumption relative to the control without 2C6MP. However CH_4_ oxidation was completely inhibited for > 20 months in the presence of 10 mM 2C6MP. In all systems, O_2_ consumption tracked with CH_4_ depletion, consistent with aerobic CH_4_ oxidation. Our results are similar to those of Megraw and Knowles [13] showing ∼60% inhibition of CH_4_ oxidation by *Methylosinus trichosporium* OB3b in liquid culture containing 4 mM 2C6MP. However, Topp and Knowles [6] reported that concentrations of 2C6MP as low as 43 µM completely inhibited CH_4_ oxidation by *M. trichosporium* OB3b. The higher concentration of 2C6MP needed for inhibition of CH_4_ oxidation in our experimental systems compared to the pure culture studies of Megraw and Knowles, and Topp and Knowles, may be due to uptake of 2C6MP by the wetland sediment, as sorption of nitrapyrin to soil has been shown to limit its effectiveness [14].

**Figure 2.**
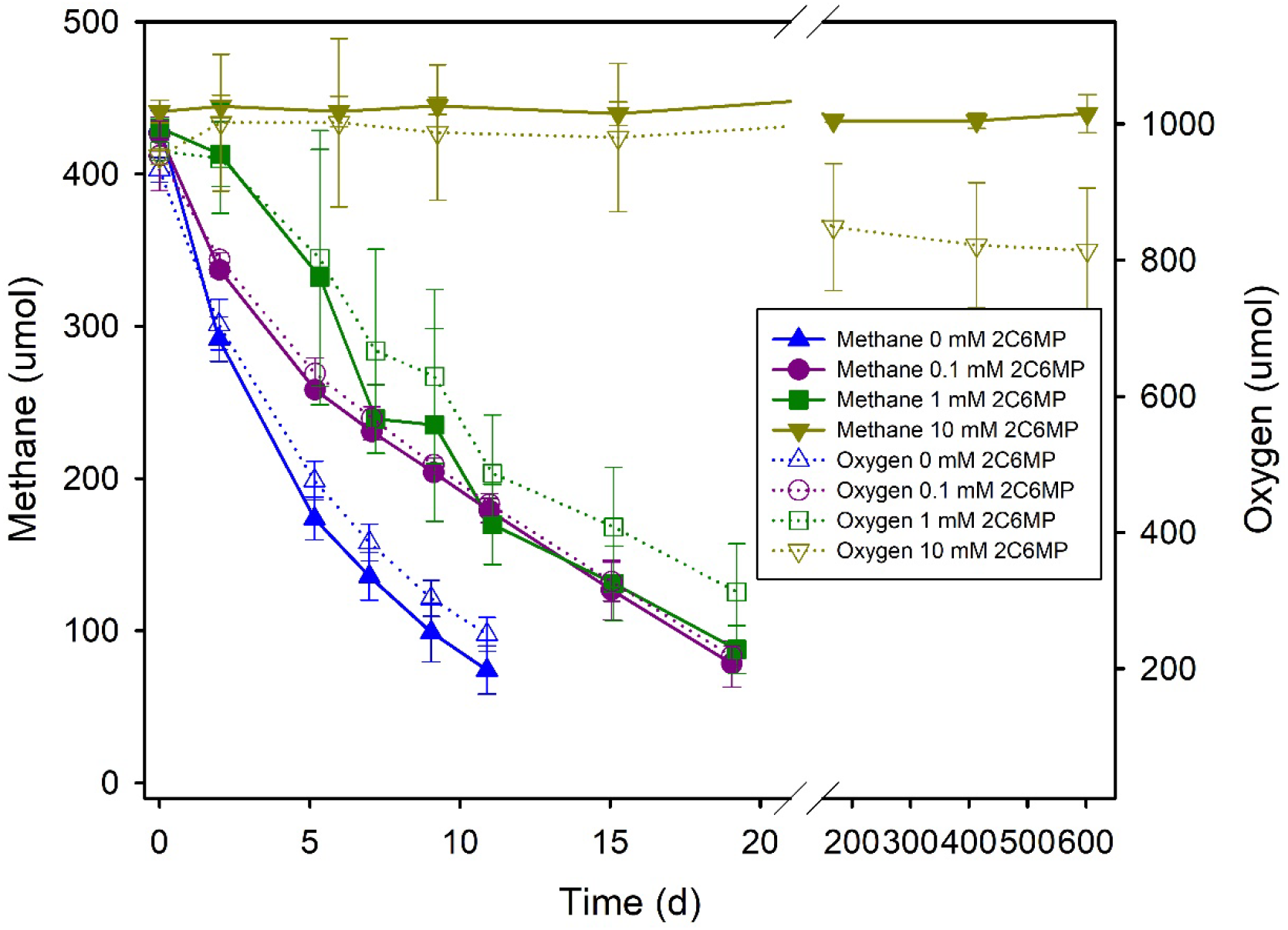
Changes in CH_4_ and O_2_ levels in the microcosm headspace over time.

Besides nitrapyrin and 2C6MP, other pyridine derivatives (as well as pyridine itself), have been studied as potential inhibitors of CH_4_ oxidation. At a concentration of 1 mM, pyridine inhibited CH_4_ oxidation by ∼60% in pure cultures of *M. trichosporium* OB3b [15]. Picolinic acid (pyridine-2-carboxylic acid) inhibited > 90% of CH_4_ oxidation by *M. trichosporium* OB3b at a concentration of 40 µM, but nicotinic acid (pyridine-3-carboxylic acid) and isonicotinic acid (pyridine-4-carboxylic acid) were largely ineffective (only 5.5% and 4.8% inhibition, respectively) [13]. These results highlight the significance of ring constituents and their location on the ring on the inhibition of microbial CH_4_ oxidation. Although relatively high concentrations of 2C6MP were needed to completely inhibit CH_4_ oxidation in our study, the effect was long lasting (> 20 months) compared to picolinic acid, which was readily degraded in soil, resulting in the loss if its inhibiotory effects within 4 days [13], and nitrapyrin, which has a reported half-life of 5–42 days in soils [14].

In addition to pyridine, pyridine derivatives, and other aromatic *N*-heterocycles (e.g., 8-hydroxyquinoline) [15,16], a diverse range of compounds have been shown to inhibit CH_4_ oxidation [17], including C_2_ hydrocarbons (e.g., ethene and acetylene) [8,18,19], halogenated aliphatic hydrocarbons (e.g., fluoromethane, difluoromethane, and 1,3-dichloropropene) [18,20-23], and dimethyl sulfoxide [24].

### 3.2. Microbial Community Dynamics

Analysis of similarity (ANOSIM) of weighted UniFrac distances between groups of triplicate samples supported a primary distinction of the inoculum relative to the enrichments (R=0.999) and a secondary distinction between microcosms containing 2C6MP versus those without (R=0.464 [0.1 mM]; R=0.894 [1 mM]) (Figure 3).

**Figure 3.**
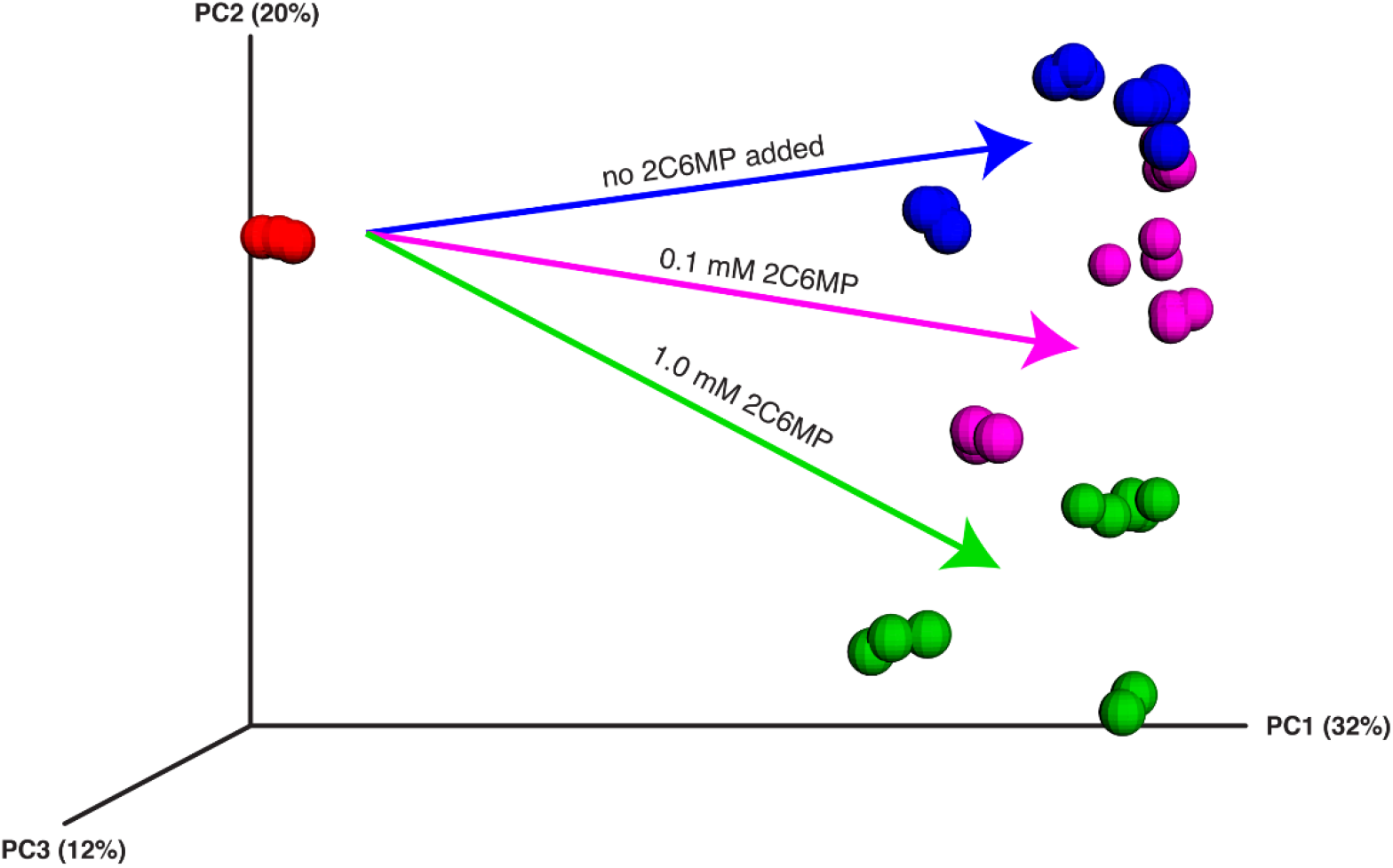
Principal coordinates analysis (PCoA) of microbial communities in Old Woman Creek inoculum and microcosms based on 16S rRNA amplicon libraries using the Illumina MiSeq platform. Distance between each data point is proportional to the weighted UniFrac distance. Triplicate DNA extractions were taken from each sample and are plotted separately.

The inoculum was dominated by members of the *Proteobacteria* (49.9%), and to a lesser extent by *Bacteroidetes* (8.8%), *Acidobacteria* (8.9%), and *Verrucomicrobia* (4.4%). In all microcosms, with or without 2C6MP, *Proteobacteria* expanded to comprise 65–70% of the total. In the absence of inhibitor, members of the *Methylococcaceae* and *Methylophilaceae* increased in relative abundance from < 0.1% in the inoculum to 10.8% and 18.1%, of the total community respectively (Figure 4). At both 0.1 and 1 mM concentrations of the inhibitor, the *Methylococcaceae* were less abundant, representing 7.9% and 8.4% respectively. No inhibition of the *Methylophilaceae* was seen at the lowest concentration of 2C6MP, but at 1 mM this taxon was only 11.4% of the total. In contrast, members of the *Crenotrichaceae*, which includes methanotrophs [25] that posess an ‘unusual’ pMMO [26], increased in relative abundance with greater amounts of inhibitor, representing 8.6% of the total at 0.1 mM and 12.3% at 1 mM, compared to only 4.1% when no inhibitor was present. Interestingly, members of *Rhodobacteraceae* were also more abundant in the presence of 0.1 (9.6%) and 1 mM (8.6%) 2C6MP than in the control (< 0.2%). Further differentiation in the relative abundance of dominant populations as a function of 2C6MP concentration is apparent at the genus level (Figure 5).

**Figure 4.**
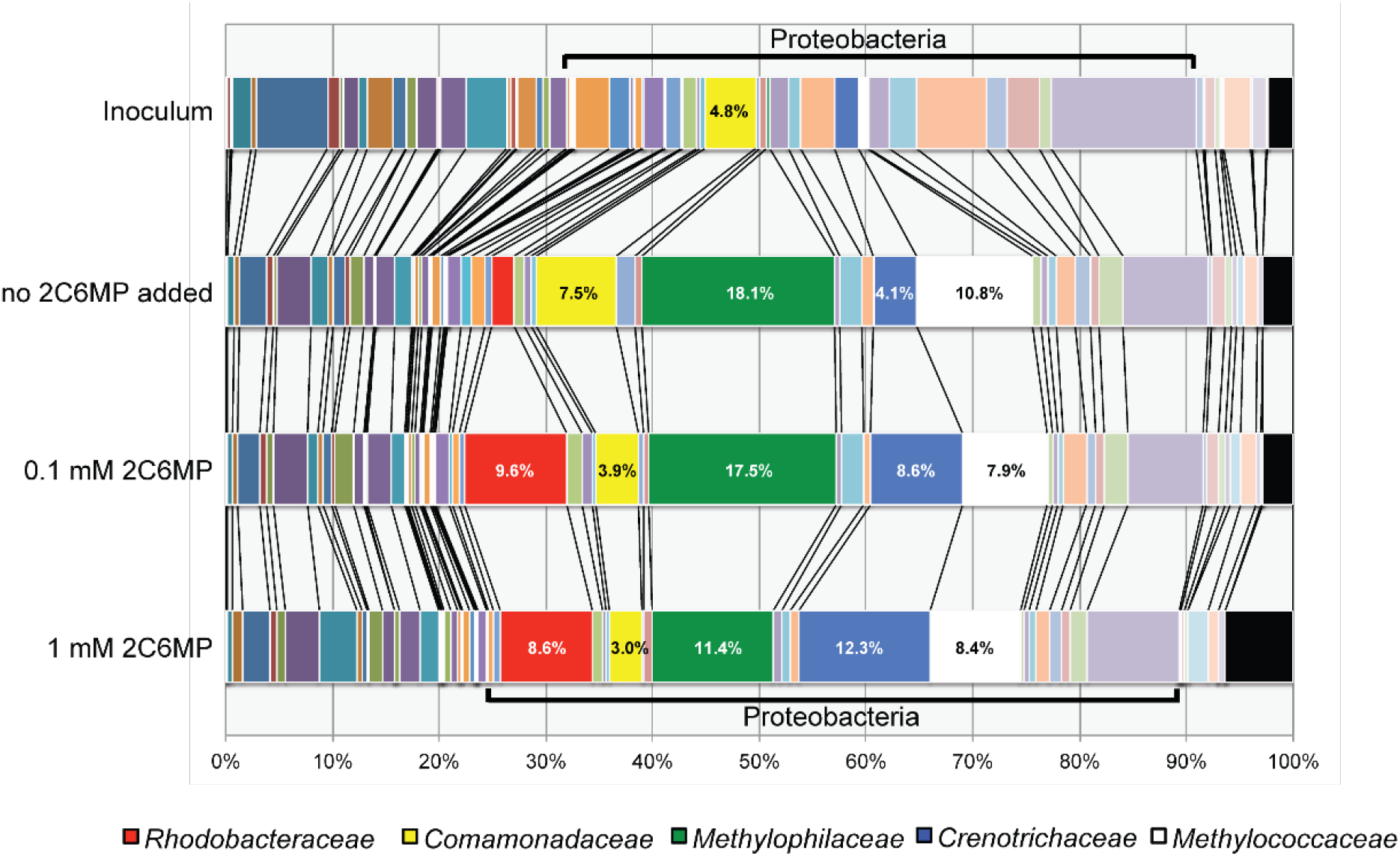
Bar graphs represent average family-level compositions for each treatment. Highlighted are key Proteobacteria families that increase in abundance relative to the inoculum.

**Figure 5.**
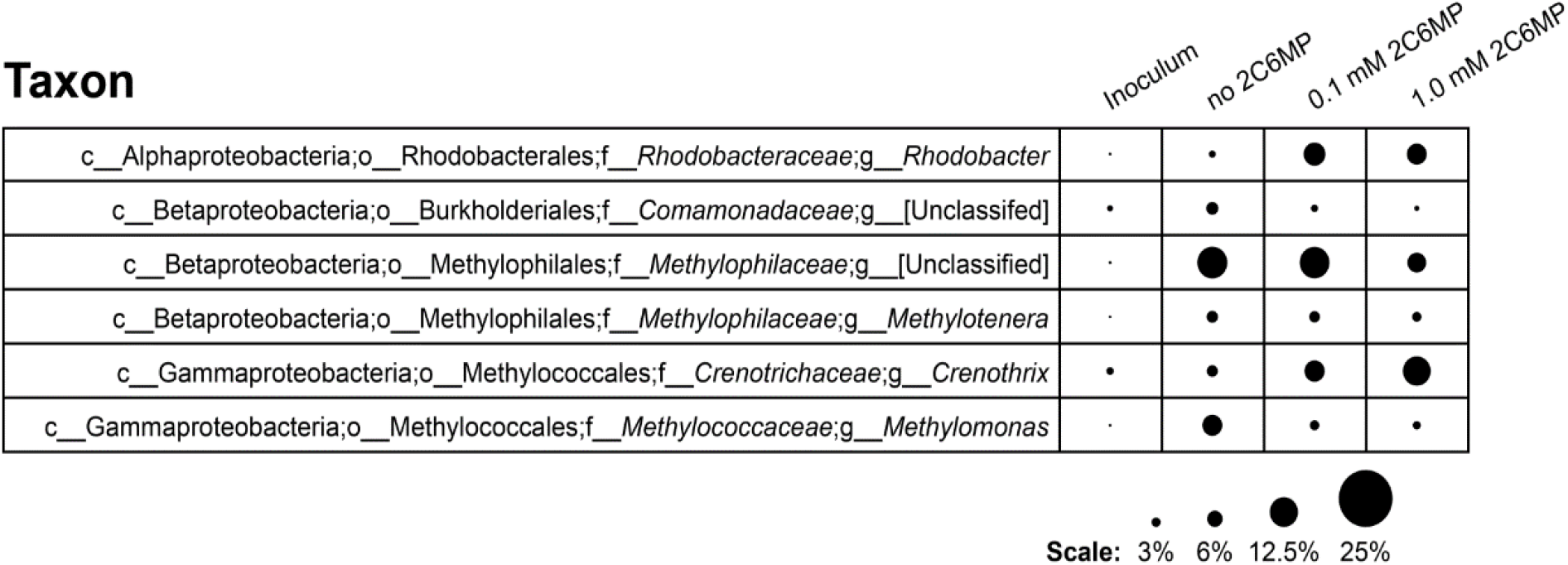
Taxonomy and average relative abundance of selected taxa in original Old Woman Creek inoculum and microcosms. Sequences were classified to the genus level of the Greengenes taxonomy using QIIME. The area of each circle is proportional to the average relative abundance of each genus.

### 3.3. Conclusions

At the highest 2C6MP concentration tested (10 mM), CH_4_ oxidation was completely inhibited for >20 months. Although lower concentrations of 2C6MP (0.1 mM and 1 mM) exhibited similar levels of inhibition of CH_4_ oxidation, there were distinct changes in the populations of putative aerobic methanotrophs relative to the amount of 2C6MP present. Though not as toxic as some other inhibitors of microbial CH_4_ oxidation (i.e., higher concentrations of 2C6MP are needed for compelte inhibiion), 2C6MP may provide long-term inhibition and may therefore be useful as a CH_4_ oxidation inibitor in laboratory studies.

## Author Contributions

Conceptualization E.J.O.; Formal analysis D.A.A and T.M.F; Funding acquisition E.J.O; Investigation E.J.O, J.C.K, and S.M.O; Resources K.K.A; Project administration E.J.O; Visualization D.A.A and T.M.F; Writing-original draft E.J.O; and Writing-review & editing E.J.O, D.A.A, K.K.A, T.M.F, J.C.K, and S.M.O. All authors have read and agreed to the published version of the manuscript.

## Acknowledgments

The authors thank the Old Woman Creek National Estuarine Research Reserve for access to the site.

